# Genome-wide SNVs revealed reticulate evolutionary history of an Eurasia-African songbird radiation

**DOI:** 10.1101/2025.08.29.673141

**Authors:** Chuan Jiang, Yang Liu, Wenwen Zhu, Nassoro Mohamed, Jiazheng Jin, Bo Li

**Affiliations:** College of Wildlife and Protected Area, Northeast Forestry University, Harbin 150040, China; School of Ecology, Sun Yat-Sen University, Shenzhen 518107, China; College of Life Science and Technology, Jinan University, Guangzhou 510632, China; College of African Wildlife Management, Mweka, Moshi P.O Box 3031, Kilimanjaro, Tanzania; BGI Research, Wuhan 430074, China; State Forestry and Grassland Administration Detecting Center of Wildlife, Harbin 150040, China

**Keywords:** Introgression, Phylogenomics, Stonechat, Single nucleotide variants

## Abstract

Phylogenomics has the power to uncover complex evolutionary hypothesis across taxa, thereby allowing important glimpses into the evolutionary history, such as incomplete lineage sorting (ILS) and introgression among closely related species. The *Saxicola torquatus* complex comprises several widely distributed insectivorous bird species across the Palearctic and Afrotropical realms. Nevertheless, the evolutionary history within this complex remains contentious, and previous findings suggest complication caused by cytonuclear discordance. Here, we conducted a phylogenomic analysis of nearly complete taxa within the complex using genome-wide SNVs to investigate potential causes of phylogenetic discordance across genomic regions and elucidate its evolutionary history. We detected multiple reticulate events during their diversification, leading to extensive phylogenetic discordance across the genome. The mitochondrial genome and W chromosome exhibited a consistent gene tree that conflicted with the species tree, with introgression being the most plausible explanation among several hypotheses. The Z chromosome was less affected by ILS/introgression and exhibited relatively stable local phylogenies compared to autosomes. Furthermore, we found that strongly introgressed regions were predominantly concentrated at the ends of smaller chromosomes, and the strength of introgression among species pairs showed consistently positive correlations, likely attributed to shared recombination hotspots in these regions. These findings stress the heterogeneous genomic landscape influenced by intricate evolutionary processes that endorse a reticulate evolution model for global speciation in the *S. torquatus* complex.

## Introduction

Early phylogenetic studies, often limited to few loci, are increasingly challenged by genome-scale data revealing pervasive phylogenetic discordance—discordance among gene trees or between gene and species trees (Edelman et al. 2019, Jiang et al. 2024, Zhang et al. 2024). Two major causes of this heterogeneity are incomplete lineage sorting (ILS) and introgression (Steenwyk et al. 2023). ILS results from the random retention of ancestral polymorphisms during population divergence, increasing with larger ancestral effective population sizes (*Ne*) and shorter internode times (Avise and Robinson 2008, Rivas-González et al. 2023). Introgression occurring through interspecific hybridization and backcrossing, where foreign alleles are incorporated into the recipient genome via recombination, typically following secondary contact between isolated populations and leading to reticulate evolution (Payseur and Rieseberg 2016, Ottenburghs et al. 2017). These two processes often act concurrently but contribute differentially across radiating lineages due to lineage-specific traits (Myers et al. 2024, Baraf et al. 2025). They not only influence the robustness of phylogenetic inference but also shape trait evolution and reflect biogeographic history (Feng et al. 2022, Hu et al. 2022, Rivas-González et al. 2023, Musher et al. 2024). Therefore, evaluating their respective contributions is critical for understanding the evolutionary history of focal clades.

Beyond lineage-specific traits, the effects of ILS and introgression also vary within lineages depending on the inheritance modes of different genomic components (Nikelski et al. 2023, Chase et al. 2024). For example, the Z chromosome, due to its hemizygosity in females, contains more incompatibility loci than autosomes, along with a reduced *Ne* and recombination rate, making it less prone to ILS and introgression (Edelman et al. 2019, Chase et al. 2024). Two maternally inherited markers, the mitogenome and the non-recombining region of the W chromosome (NRW), show stronger effects of hemizygosity and lower *Ne* than the Z chromosome and lack recombination, further minimizing ILS and introgression (Moore 1995, Smeds et al. 2015, Mackiewicz et al. 2019). Their strict maternal inheritance, however, creates a distinct dynamic: after hybridization, they may persist in the recipient population and rapidly reach fixation due to their small *Ne*, resulting in phylogenetic discordance with the species tree (e.g., mitogenome capture) (Toews and Brelsford 2012, Nikelski et al. 2023). Moreover, due to the functional coupling between mitochondrial and nuclear genomes (Shi et al. 2025), introgressed mitogenomes may induce mitonuclear incompatibilities (Sloan et al. 2017), driving the co-introgression of mitonuclear genes alongside the mitogenome to maintain mitochondrial function (Hill 2019, Nikelski et al. 2023). Analyzing these phenomena not only facilitates accurate reconstruction of species tree but also highlights the differential roles of genomic components during lineage radiations.

Due to the widespread occurrence of avian hybridization (Ottenburghs et al. 2015, Ottenburghs 2023) and the slow evolution of intrinsic postzygotic isolation (Fitzpatrick 2004), introgression frequently occurs in birds. Introgression plays a dual role in speciation (Musher et al. 2024): on one hand, it can introduce beneficial standing genetic variation from other populations, facilitating adaptation to new environments (Meier et al. 2019, Hu et al. 2022); on the other hand, it may disrupt reproductive isolation barriers between species, thereby hindering speciation (Taylor et al. 2006, Webb et al. 2011). The resulting introgression landscape—that is, the heterogeneous acceptance of foreign genetic material across the genome—may be shaped by multiple factors, including natural selection strength, local recombination rates, and the distribution of reproductive isolation genes (Gante et al. 2016, Ravinet et al. 2017, Martin and Jiggins 2017, Moran et al. 2021, Veller et al. 2023, Musher et al. 2024, Blain et al. 2025). However, current knowledge about the evolutionary role of introgression and its underlying drivers remains limited to a few taxa. Expanding research to include more non-model species will enhance our understanding of the genetic mechanisms underlying speciation and species maintenance.

The *Saxicola torquatus* complex comprises several widely distributed insectivorous bird species across the Palaearctic and Afrotropical realms, with a long-debated taxonomic history (Zink et al. 2009, Opaev et al. 2018). Historically, this complex was considered a superspecies, the common stonechat *S. torquatus*, encompassing multiple subspecies (Zink et al. 2009, Svensson et al. 2012, Opaev et al. 2018). Based on morphology, vocalizations, habitat preference, mitochondrial genes, and genomic data from a limited number of taxa (Wittmann et al. 1995, Urquhart 2002, Illera et al. 2008, Zink et al. 2009, Svensson et al. 2012, Hellström and Norevik 2014, Van Doren et al. 2017, Opaev et al. 2018, Loskot and Bakhtadze 2020, Hellström and Waern 2023, Zhao et al. 2023, Jiang et al. 2025), the *S. torquatus* complex is now tentatively classified into six species: *S. torquatus*, *S. tectes*, *S. rubicola*, *S. dacotiae*, *S. maurus*, and *S. stejnegeri* (Winkler et al. 2020, Gill et al. 2025). However, the species-level status of *S. stejnegeri* remains debated due to its parapatric distribution with *S. maurus* and potential hybridization (Hellström and Norevik 2014, Hellström and Waern 2023), leading to the suggestion that it might be a subspecies of *S. maurus*, namely *S. m. stejnegeri* (Chesser et al. 2022, Gill et al. 2025, AviList Core Team 2025). This ambiguity is partly attributed to previous studies on this species relying on mitochondrial genes, which have not yielded a conclusive phylogeny. Additionally, the phylogenetic position of *S. maurus* shows cytonuclear discordance (Figure S1), likely due to ILS, ancient introgression, or other biological processes (Zink et al. 2009, Van Doren et al. 2017), underscoring the complexity of their evolutionary history. Thus, the *Saxicola torquatus* complex represents a promising system for studying how ILS and introgression shape gene-species tree conflicts.

In this study, we performed whole-genome sequencing on the key lineage *S. stejnegeri* within the *S. torquatus* complex and integrated it with published genomic data from several other lineages within this complex. This ultimately covers nearly complete taxa lineages within the *S. torquatus* complex (except *S. tectes*), as well as *S. rubetra* outside the complex. To resolve their evolutionary history, we first reconstructed the phylogeny of this complex using different genomic regions and assessed the levels of ILS and introgression. Based on the discovered reticulate evolution, we further analyzed the introgression landscape and correlations between four species pairs and performed functional enrichment analysis of introgressed genes to explore potential drivers of introgression patterns. We also conducted a mitonuclear co-introgression test for an ancient hybridization event and discussed the causes of cytonuclear discordance in the phylogenetic position of *S. maurus*. Additionally, we reconstructed the demographic history over the past million years of four species with individual sequencing data. Finally, based on the reconstructed phylogeny and hybridization history, we provided new insights into the taxonomic classification and colonization history of the *S. torquatus* complex.

## Materials and methods

### Sample Collection and Sequencing

The *S. stejnegeri* carcass sample was obtained from Bei’an City, Heilongjiang Province, seized by the State Forestry and Grassland Administration Wildlife Detection Center (Harbin, China) in an illegal capture case. Total genomic DNA was extracted from muscle tissue using a tissue extraction kit (UElandy, Suzhou, China). High-quality DNA was fragmented by sonication, and a library with an insert size of ∼350 bp was prepared on the DNBSEQ platform following the manufacturer’s instructions (MGIEasy Universal DNA Library Preparation Kit, BGI). Sequencing was performed using a 150 bp paired-end strategy on the DNBSEQ-T7 (MGI, China).

### Data Processing

In addition to the *S. stejnegeri* sequences we generated, we obtained individual sequencing data for *S. rubetra*, *S. rubicola*, and *S. rubicola*, along with pooled sequencing data for *S. torquatus* and *S. dacotiae* from the SRA database (Van Doren et al. 2017, Bergeron et al. 2023) (Table S1). All data were filtered using fastp (Chen 2023) with default parameters, and for the stlFR data of *S. rubetra* and *S. rubicola*, barcode sequences at the 3’ end of the reads were trimmed.

Quality-controlled reads were aligned to the chromosomal-level genome of male *Oenanthe melanoleuca* (GCA_029582105.1) (Peona et al. 2023) using BWA-MEM v0.7.18 (Jung and Han 2022) with default parameters. Reads with a mapping quality (MAPQ) >20 were retained and were sorted by genomic coordinates using SAMtools v1.21 (Danecek et al. 2021). PCR duplicates were removed using Picard v3.1.1 (McKenna et al. 2010). Alignment rates and coverage for each sample were calculated using the flagstat and coverage modules in SAMtools, respectively. Additionally, sex was determined by the normalized Z chromosome depth (Z depth / mean autosomal depth), with expected values of ∼1 for males and ∼0.5 for females.

Based on the deduplicated mapping results, single nucleotide variants (SNVs) were called using HaplotypeCaller, and joint genotyping was performed on all samples using GenotypeGVCFs in GATK v4.5 (McKenna et al. 2010). Next, Indels were excluded using the SelectVariants module, and hard-filtering was applied with the VariantFiltration module in GATK using the following criteria: RPRS < -8, SOR > 3.0, QD < 2.0, FS > 60.0, MQ < 40.0, and MQRankSum < −12.5. Finally, SNVs were soft-filtered using Bcftools v1.20 (Danecek et al. 2021) and Vcftools v 0.1.16 (Danecek et al. 2011), retaining only biallelic loci, excluding SNVs missing in any sample, and removing heterozygous sites on the Z chromosome in any female and pooled samples. Additionally, we calculated both the genome-wide autosomal heterozygosity rate and the heterozygosity rates in 100 kb non-overlapping windows using Vcftools v0.1.16 and Bedtools v2.31.1 (Quinlan and Hall 2010).

### Phylogenetic analysis of concatenated whole-genome SNV

Based on the high-quality SNVs obtained above, we first constructed species trees using the connected SNVs. Three main approaches were applied: (1) maximum likelihood (ML) methods implemented in RAxML-NG v1.2.2 (Kozlov et al. 2019) and IQTREE v2.3.6 (Minh et al. 2020); (2) site-based coalescent methods using quartet topology frequencies, implemented in CASTER v1.19.1.4 (Zhang et al. 2025a) and PAUP v4.0 (SVDquartets) (Chifman and Kubatko 2014); and (3) the Bayesian multispecies coalescent method SNAPP, implemented in BEAST v2.6.3 (Bryant et al. 2012).

In RAxML-NG, the GTR+Γ nucleotide substitution model was used, with 1,000 bootstrap replicates. In IQTREE, we used the GTR+ASC model to correct for SNP ascertainment bias, also performing 1,000 bootstrap replicates. In CASTER, species trees were constructed based on individual sites patterns (caster-site) and pairwise site patterns (caster-pair), respectively.

For methods requiring unlinked loci (SVDquartets and SNAPP), we used VCFtools v0.1.16 to thin the dataset, ensuring a minimum inter-SNP distance of 1,000 bp. In SVDquartets, all possible quartets were evaluated, and 1000 bootstrap replicates were performed. SNAPP was run for 10 million generations with sampling every 500 generations. Convergence was assessed using Tracer v1.7 (Rambaut et al. 2018), in which the first 25% of trees were discarded as burn-in and all parameters were confirmed to have effective sample sizes greater than 200. Posterior SNAPP trees were visualized in DensiTree v2.2.7 (Bouckaert 2010).

### Phylogenetic reconstruction of window-based strategy

We further performed species tree inference using a window-based strategy. First, we constructed genome fragment (GF) trees along the autosomes and Z chromosome with 50 kb non-overlapping windows using genomics_general (https://github.com/simonhmartin/genomics_general). For each GF tree, a ML tree was inferred under the GTR+Γ substitution model using RAxML-NG v1.2.2, with 100 bootstrap replicates. The species tree was then obtained from the GF trees using Astral-hybrid v1.19.3.7 (Zhang and Mirarab 2022), based on node support weights under default parameters.

To examine phylogenetic discordance across the genome, TWISST (Martin and Van Belleghem 2017) was used to compute the proportions and genomic distribution of alternative topologies across all GF trees. The GF trees were visualized using DensiTree v2.2.7, with a randomly selected subset of 1,500 trees.

### Phylogenetic analysis of NRW and mitogenome

To examine the phylogeny of NRW, we downloaded the NRW sequences of *Ficedula albicollis* (GCA_900067835.1) (Smeds et al. 2015) and selected all female samples and pooled samples (Table S1) to obtain NRW SNVs using the same method as described previously. To exclude potential effects of mapping errors and homologous sequences from the Z chromosome,we retained only reads with a mapping quality of 60 in BWA mapping and kept only SNV sites that were homozygous across all samples during the soft filtering. We then constructed a ML tree for NRW using RAxML-NG v1.2.2 under the GTR+Γ substitution model, with 1000 bootstrap replicates.

For the mitochondrial (mt) genome (mitogenome), we first assembled the 13 protein-coding genes (PCGs) of seven species using MitoFinder v1.4.1 (Allio et al. 2020) and then aligned them with MAFFT v7.313 (Katoh and Standley 2013). The ML tree of the 13 PCGs was subsequently constructed using RAxML-NG v1.2.2 under the GTR+Γ substitution model, with 1000 bootstrap replicates.

### Hybridization detection

We first calculated the frequencies of three topologies per node from genome-wide GF trees using DiscoVista (Zhang et al. 2018), comparing the proportions of two alternative topologies to assess introgression and ILS. We then applied QuIBL to evaluate introgression/ILS levels using branch length information from GF trees (Edelman et al. 2019). Due to its sensitivity to recombination, we extracted a window every 500 kb to ensure the independence between windows for analysis. Each species triplet was analyzed under both the ‘ILS-only’ and ‘ILS+introgression’ models. Following recommendations, introgression is considered only when the BIC of the ’ILS+introgression’ model is at least 10 units smaller than the Bayesian information criterion (BIC) of the ’ILS-only’ model; otherwise, only ILS is present. The results reflecting introgression and/or ILS between species pairs were averaged across all relevant triplets, considering only two alternative topologies.

We further assessed introgression using a site-based approach. First, we calculated Patterson’s *D* statistics (ABBA-BABA test) and *f*_4_-ratio using Dsuite v0.5 (The Heliconius Genome Consortium 2012, Malinsky et al. 2021). We fixed *O. melanoleuca* as the outgroup and computed *D*-values/*f*_4_-ratio for all possible combinations based on the species tree topology. We applied a standard block-jackknife procedure to assess the statistical significance of the *D* statistics and *f*_4_-ratio by dividing the dataset into 200 Jackknife blocks. Only combinations with a Z-score > 3 were retained, and the heatmap of *D* statistic and *f*_4_- ratio was generated using the Ruby script plot_d.rb (https://github.com/mmatschiner/tutorials/tree/master/analysis_of_introgression_with_snp_da ta). Since different combinations were non-independent due to shared branches on the phylogenetic tree, this may result in highly correlated values when ancient gene flow is present. We therefore utilized the Fbranch module in Dsuite to calculate branch-specific estimates of gene flow (f-branch statistics, *f_b_*), which can indicate introgression between extant species and ancestral lineages. The *f_b_* statistic values for each branch were calculated from *f*_4_-ratio values that passed the significance threshold (Z > 3), and were visualized using the dtools.py script.

We inferred the phylogenetic network using SNaQ implemented in the Julia package PhyloNetworks v0.16.2 (Solís-Lemus et al. 2017), based on the same set of GF trees used for QuIBL analysis. The search allowed for up to four hybridization events (h = 4), initialized with the species tree as the starting topology. For subsequent optimizations, we used the network estimated in the previous step as the starting network. The optimal number of reticulation events was determined by identifying the inflection point showing the steepest improvement in log-pseudolikelihood scores. The optimal number of reticulation events was determined by identifying the inflection point showing the steepest improvement in log-pseudolikelihood scores. Following model selection, we ran 100 bootstrap replicates based on the 100 bootstrap trees generated from each GF tree, employing default settings. The phylogenetic network is also inferred using the ‘InferNetwork_MPL’ command in PhyloNet v3.8.2 (Yu et al. 2014), allowing for up to four reticulation events and starting from the species tree topology. For each reticulation setting, 100 independent searches are performed to avoid local optima. The optimal network is selected based on the highest log-pseudolikelihood score.

Gene flow was also inferred by utilizing allele frequency information from different species using TreeMix v1.13 (Pickrell and Pritchard 2012). The same SNV dataset of unlinked loci as in the SANPP analysis was used, with outgroups further removed. First, a set of simulations was performed to define the best migration edges, where migration edges were set from 0 to 7, each with 10 parallel runs. The outputs were then analyzed using the R package OptM (Fitak 2021), and the optimal number of migration edges was selected based on the second-order rate of change in likelihood (Δm).

### Introgression pattern and introgressed gene enrichment analysis

We investigated the genomic landscape of several hybridization events using the ABBA-BABA test with 50 kb non-overlapping windows, implemented in genomics_general. Four species combinations were used to examine four specific hybridization events, with *O. melanoleuca* fixed as the outgroup: ((*S. torquatus*, *S. rubicola*) *S. maurus*) to examine introgression between *S. maurus* and *S. rubicola*; ((*S. torquatus*, *S. dacotiae*) *S. maurus*) for *S. maurus* and *S. dacotiae*; ((*S. torquatus*, *S. maurus*) *S. stejnegeri*) for *S. maurus* and *S. stejnegeri*; and ((*S. rubicola*, *S. dacotiae*) *S. torquatus*) for *S. dacotiae* and *S. torquatus*. These combinations were chosen to capture strongest signals of introgression among all possible triads while minimizing potential gene flow between P1 and P2. Due to the sensitivity of the *D* statistic to genomic diversity in window-based analyses, we used *f*_d_, a modified *D* statistic (Martin et al. 2015), as the introgression measure. Finally, we performed Spearman rank correlation analyses to evaluate the similarity of introgressed regions across species pairs. As negative *f*_d_ values are biologically meaningless, we retained only positive *f*_d_ values and truncated any values exceeding 1.

We further examined whether introgressed genes between each species pair were significantly enriched for specific functions. Based on the same combinations described above, we calculated the *f*_d_ values for 15,782 PCGs located on the autosomes and the Z chromosome for each species pair. Genes with *f*_d_ values greater than 0.2 were considered candidate introgressed genes and were subjected to Gene Ontology (GO) and Kyoto Encyclopedia of Genes and Genomes (KEGG) enrichment analyses using clusterProfiler v4.16.0 (Xu et al. 2024).

### Test for mitonuclear co-introgression

Based on the observed mitonuclear discordance within the *S. torquatus* complex, we propose a hypothesis that this conflict arose from mitogenome capture by *S. maurus* from either the most recent common ancestor (MRCA) of *S. rubicola* and *S. dacotiae* or a sister species to this ancestor—which may represent a cryptic, unsampled lineage or an extinct ghost lineage (hereafter ‘unknown species’). Given the coevolution between mitochondrial and nuclear genomes (Shi et al. 2025), we predict that mitonuclear genes will exhibit co-introgression with the mitogenome due to selection pressures driven by mitonuclear incompatibilities—if mitogenome capture is present. Since the genetic material of the unknown species is unavailable, we use *S. rubicola* and *S. dacotiae* as proxies for the unknown species in our analyses, as they share similar genetic components with it, although this approach may lead to false negatives.

We compiled a list of mitonuclear genes by consulting various resources on mitochondrial and mitonuclear dynamics, and prior similar work(Diodato et al. 2014, Greber and Ban 2016, Nikelski et al. 2023). These included only PCGs whose products directly interact with mtDNA or its immediate translation products. After removing genes that were not annotated or lacked specific genomic locations in the reference genome, 202 mitonuclear genes remained (Table S2). The remaining 15,580 PCGs were classified as non-mitonuclear.

We used a gene-based approach to calculate the introgression signal strength for each gene and applied a threshold to categorize them as introgressed or non-introgressed. We employed chi-square and Fisher’s exact tests to determine whether mitonuclear genes were over-representation among the introgressed genes. This analysis was repeated across different introgression indicators (*D*, *f*_d_, and *f*_dM_), thresholds levels (top 1%, 5%, 10%, 15%, 20%, and 25%), and species combinations (Table S3). In all ABBA-BABA test configurations (((P1, P2), P3), Outgroup), *S. torquatus* was fixed as P1 and *S. maurus* as P3, while P2 and the outgroup varied.

Since gene-based methods may introduce computational randomness due to the short length of some genes, and because regulatory elements near genes could also co-introgress adaptively with mitonuclear genes, we repeated the analysis using a window-based approach. Specifically, we calculated introgression signal strength for each 50 kb non-overlapping window and used a threshold to classify the windows as either introgressed or non-introgressed. We assigned each gene to these windows by minimizing the absolute difference between the gene center location and the window center location, thereby categorizing genes as introgressed or non-introgressed. The remaining steps were consistent with the gene-based approach (Table S4).

### Demographic History

The pairwise sequentially Markovian coalescent (PSMC) analysis was performed on genomic data from four individual sequencing data (Table S1) to infer their demographic history. Initially, diploid consensus sequence was generated for each individual using SAMtools v1.21 and BCFtools v1.20 based on the previously obtained mapping results, with sites where sequencing depth was below one-third or above twice the average depth masked. PSMC v0.6.5 (Li and Durbin 2011) was run with the parameters “–N25 –t15 –5 –p 4+25*2+4+6”. To assess the robustness of the results, 100 bootstrap replicates were performed by randomly sampling 5-Mb sequence segments from the consensus genome sequence with replacement. The generation time for all species was set to 2 years, and the mutation rate was set at 0.33% per million years (Zhang et al. 2014).

## Result

### Sex determination and variant discovery

We generated ∼33 Gb of data from whole-genome sequencing of *S. stejnegeri* (Table S1). Combined with downloaded data from five *Saxicola* species and one outgroup, the total dataset reached ∼668G (∼87X mean depth). After quality filtering, ∼17.27% ( ∼115G ) of the bases were trimmed due to low quality, complexity, or adapter contamination. Mapping to the reference showed high-quality alignment rate (97.4% to 99.08%), except for *S. torquatus* (59.1%), likely due to bacterial contamination, as noted by the data releasers (Van Doren et al. 2017). Because these bacterial reads were unmapped, they do not affect downstream analyses. Alignment coverage ranged from 95.9% to 99.4%, indicating high completeness. Gender identification based on the normalized Z-depth indicated that only *S. stejnegeri* was male (Figure S2). Two pooled samples (*S. torquatus* and *S. dacotiae*) showed intermediate normalized Z-depth values (0.71 and 0.79, respectively), matching their sex ratios (male .3: female = 18:33 and 38:18).

Following SNV filtering, a total of 45,784,604 high-quality SNVs were identified. Species-specific SNVs were most abundant in the outgroup (15,636,339) and *S. rubetra* (12,174,382), while the five *S. torquatus complex* species had 1,226,091 to 3,333,538 (Table S1). Genome-wide heterozygosity varied considerably among species, ranging from 0.017 to 0.198, which may partially reflect different demographic histories (Table S1, Figure S3). Notably, the two pooled samples showed moderate-to-low heterozygosity (*S. dacotiae* = 0.017, *S. torquatus* = 0.058), with distributions similar to other species, indicating their suitability for downstream analysis as individual representatives.

### Phylogenetic discordance across genome

All methods based on concatenated autosomal and Z-chromosome SNVs (45,784,604 SNVs) and the window-based strategy (20,138 windows) yielded consistent species tree topologies with maximal nodal support (Figure 1, Figure S4). The topology revealed a nested structure: *S. dacotiae* and *S. rubicola* were sister taxa, followed sequentially by *S. torquatus*, *S. maurus*, *S. stejnegeri*, and *S. rubetra*. In contrast, the mitogenome phylogeny placed *S. maurus* as sister to *S. dacotiae* and *S. rubicola*, with *S. torquatus* outside, also with full support, indicating cytonuclear discordance. Interestingly, the NRW phylogeny (71,321 SNVs based on all female individuals) mirrored the mitogenome topology, suggesting co-inheritance of the mitogenome and W chromosome. However, the interbranch lengths in the NRW ML tree were relatively short, possibly due to excessive filtering of SNVs.

**Figure 1.**
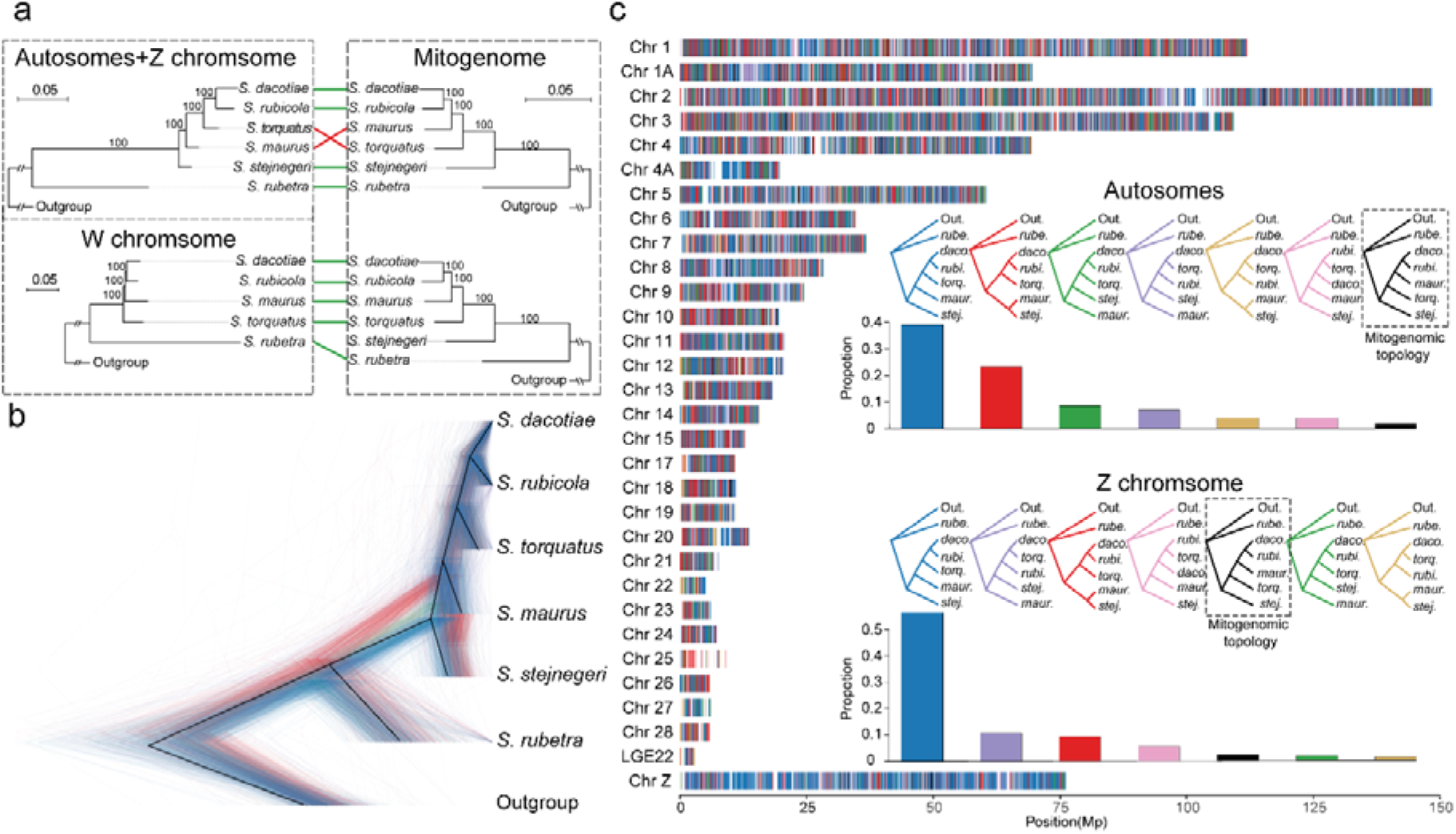
Phylogenies across different genomic regions. (a) Phylogenetic trees of the Autosomes, Z chromosome, W chromosome, and mitogenome constructed using the ML method in RAxML-NG. Solid lines connect the same taxa across trees, with green indicating phylogenetic congruence and red indicating incongruence across different trees. (b) DensiTree visualization of 1,500 sampled GF trees across the genome. Different colors represent distinct tree topologies. (c) The distribution and proportions of the seven main tree topologies across the autosomes and Z chromosome based on a 50 kb sliding window. Alternating colors along each chromosome represent different tree topologies, with colors matching those shown in the panel. White regions represent tree topologies outside the seven main topologies. The black topology, outlined by a dashed black box, corresponds to the mitogenome tree. The histograms show the proportions of each tree topology on autosomes and the Z chromosome. The abbreviations *rube.*, *daco.*, *rubi.*, *torq.*, *maur.*, *stej.* represent *S. rubetra*, *S. dacotiae*, *S. rubicola*, *S. torquatus*, *S. maurus*, and *S. stejnegeri*, respectively. The vertical axis shows the chromosome number, while the horizontal axis represents the chromosome length scaled in million base pairs (Mb).

Phylogenies reconstructed across genomic windows revealed extensive phylogenetic discordance, failing to resolve the phylogeny as a simple bifurcating tree (Figure 1 b, c). Among 20,138 GF trees, seven main topologies accounted for ∼86.9% of the total. As expected, the topology concordant with the species tree inferred from autosomal and Z-chromosome SNVs (blue topology, Figure 1 c) was most frequent, comprising ∼40.3% of trees. This topology was especially prevalent on the Z chromosome (∼56.5%), exceeding its frequency on autosomes (∼37.1%) by ∼19.4 percentage points. In contrast, the topology consistent with the mitogenome and NRW tree (black topology, Figure 1 c) was rare, representing only ∼1.7% overall (∼1.6% autosomes; ∼2.3% Z chromosome).

### Reticulate evolution within the *S. torquatus* complex

To elucidate the causes of phylogenetic discordance, we applied DiscoVista and QuIBL to distinguish between introgression and ILS, two major biological sources of discordance (Steenwyk et al. 2023). DiscoVista revealed substantial topological conflict at nodes 2 and 4, with one alternative quartet topology markedly more frequent, implicating introgression— specifically between *S. stejnegeri* and *S. maurus*, and *S. torquatus* and *S. dacotiae*—as the primary source of discordance (Figure 2 a). QuIBL analysis of 2,028 GF trees from independent windows corroborated this inference, indicating that introgression accounted for most of discordance (proportions of 0.3 and 0.11 for the two introgression events), while ILS played a minor role, peaking at 0.08 between *S. torquatus* and *S. rubicola* (Figure 2 b). *D* statistics and *f*_4_-ratio further supported these two introgression events and detected additional, moderate introgression between *S. maurus* and *S. rubicola*, and weak introgression between *S. maurus* and *S. dacotiae* (Figure 2 c). Notably, the summarized *f*_4_-ratio on shared branches (*f_b_*) revealed introgression between *S. maurus* and MRCA of *S. rubicola* and *S. dacotiae* (Figure 2 d). This may reflect a hypothesis that the cytonuclear conflict of *S. maurus* may result from mitogenome capture from an unknown species (see Materials and Methods).

**Figure 2.**
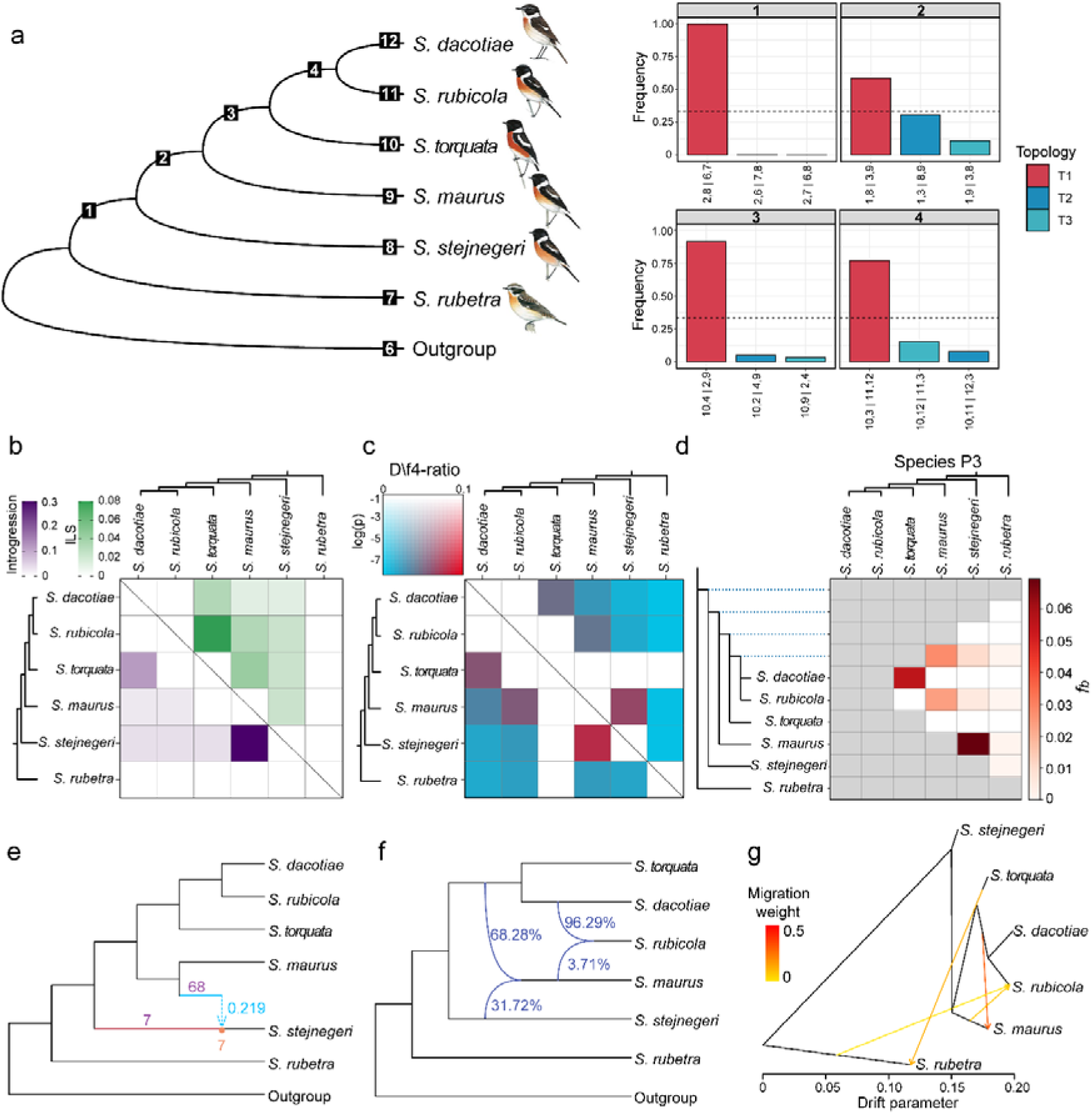
Topologies distribution per node and interspecific introgression. (a) Species tree of six *Saxicola* species (left) and the frequency of three topologies per node, calculated by DiscoVista (right). Main quartet topologies are in red, alternatives in blue. Dotted lines represent the 1/3 threshold. (b) Species pairwise metrics for the proportions of introgression (below diagonal) and ILS (above diagonal) estimated by QuIBL. Darker purple and green indicate higher proportions of introgression and ILS, respectively. (c) Species pairwise metrics show Patterson’s *D* (below diagonal) and *f*_4_-ratio (above diagonal) inferred from every trio of the six species. The color legend corresponds to the *P* value (block-jackknife) and the magnitude of introgression (*D* and *f*_4_-tario). (d) The *f_b_*statistic identifies possible introgression events from the branch of the tree on the y axis to the species on the x axis. (e) Inferred species networks using SNaQ with one hybridization event. Bootstrap values for hybrid nodes and hybrid edges are shown in orange and purple, respectively. Light blue dashed lines with numbers denote minor edges and their inheritance probabilities. (f) Inferred species networks using Phylonet with a maximum of 2 reticulations. Blue percentages show inheritance probabilities. (g) TreeMix result when 4 migration edges were allowed. Migration edge color indicates migration weight.

**Figure 3.**
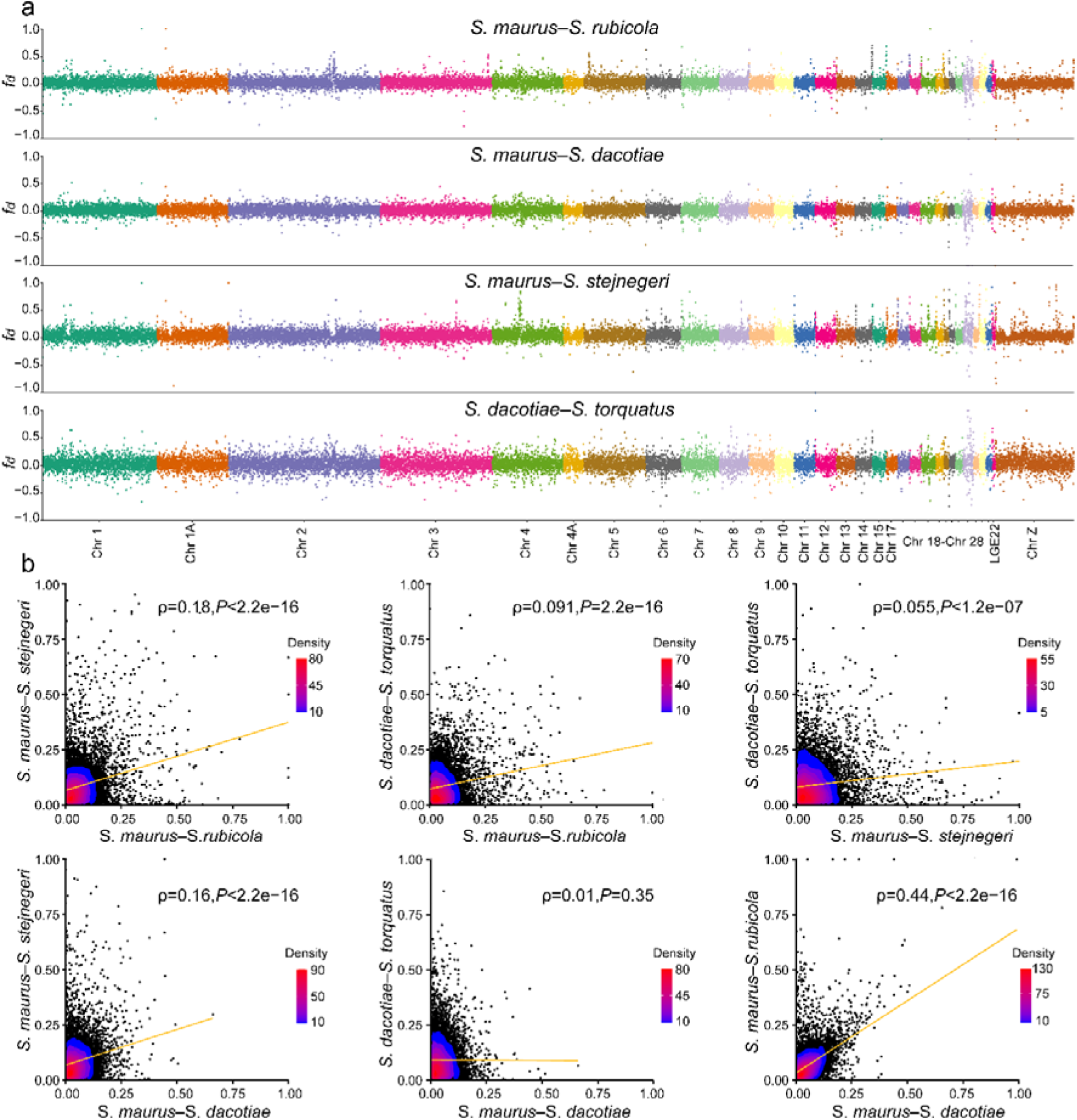
The introgression landscape of four species pairs and their pairwise correlations. (a) The introgression landscape of the four species pairs revealed by *f*_d_ calculated based on the ABBA test. Alternating colors indicate different chromosomes corresponding to the x-axis. The y-axis represents the *f*_d_ value, with values greater than 0 indicating introgression between the target two species, while values less than 0 have no biological significance. (b) Pairwise correlation of *f*_d_ among the four species pairs. Each point in the scatterplots represents one 50 Kb genomic window. Orange lines are best-fit lines, and the Spearman’s rank correlation rho (ρ) coefficient and *P* value are provided. The gradient from blue to red indicates increasing point density.

**Figure 4.**
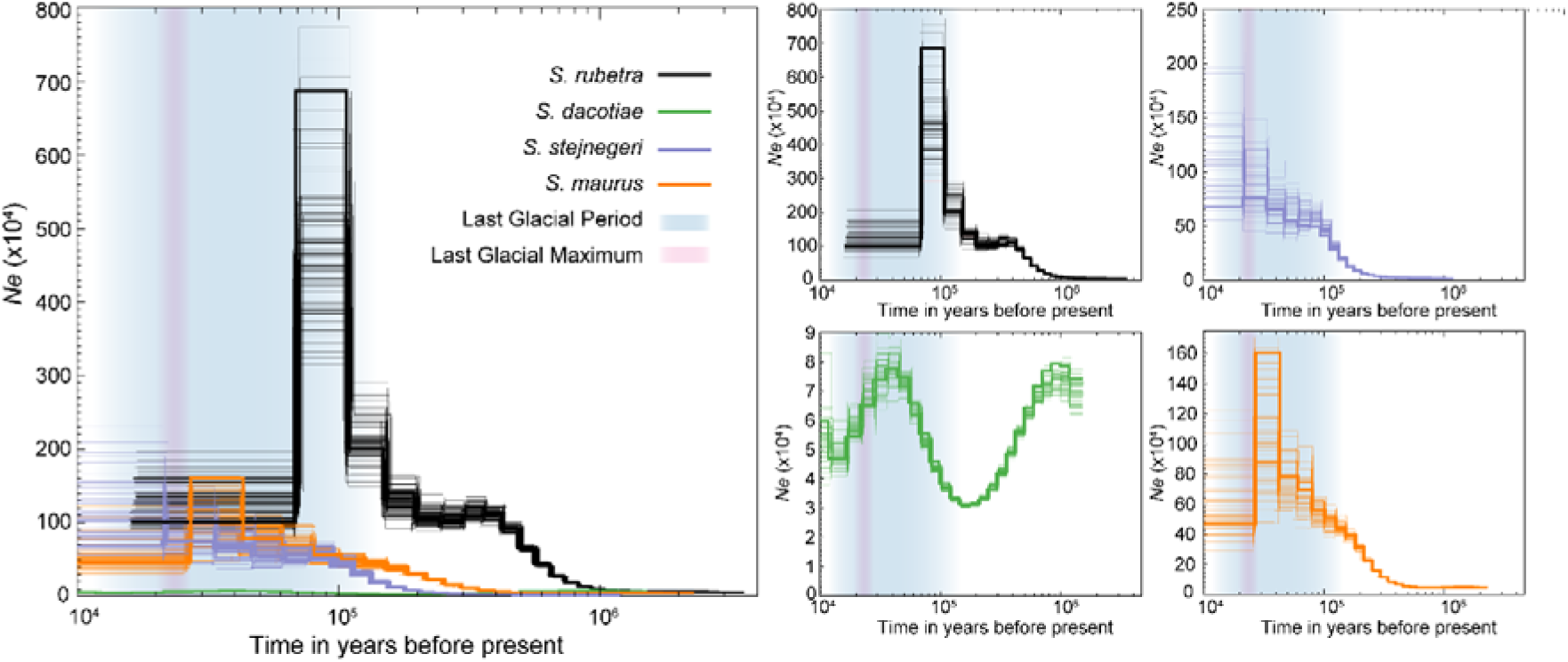
Demographic history over the past ∼1 million years for four species with individual sequencing data, inferred by PSMC. The large panel on the left shows *Ne* changes for all four species scaled to the same y-axis (*Ne*) for better comparison. The four panels on the right display the *Ne* changes for each species individually scaled to better illustrate their respective trends. The thick black, green, purple, and orange lines represent the demographic trajectories of *S. rubetra*, *S. dacotiae*, *S. stejnegeri*, and *S. maurus*, respectively, while the corresponding thin lines indicate the bootstrap analysis for each species. The blue background denotes the Last Glacial Period, and the red background represents the Last Glacial Maximum.

The SNaQ species network inference identified one optimal hybridization event (Figure S5 a), with 21.9% inheritance probability from *S. maurus* to *S. stejnegeri*. However, bootstrap support was weak: the hybrid node appeared in only 7 replicates, and the major and minor hybrid edges were supported in 68 and 7 replicates, respectively, indicating much uncertainty in introgression direction and placement. PhyloNet results showed the best fit with two reticulation events (Figure S5 b), revealing not only hybridization between *S. maurus* and *S. stejnegeri* but also between *S. maurus* and *S. rubicola*, with inheritance probabilities of 31.72% and 3.71%, respectively.

The OptM results indicated that m = 4 was the best number of migration edges in Treemix (Figure S5 c). Although Treemix did not recover the strongest introgression between *S. maurus* and *S. stejnegeri*, it revealed a notable migration edge from the MRCA of *S. rubicola* and *S. dacotiae* to *S. maurus*, which had the highest migration weight and was consistent with the results from *f_b_*. Another migration edge points from *S. maurus* to *S. rubicola*, which was also detected in the *D*-statistic, *f*_4_-ratio, *f_b_*, and PhyloNet results. However, the remaining two migration edges were not detected by other methods, including introgression from *S. torquatus* to *S. rubetra* and from *S. rubetra* to *S. rubicola*.

### Introgression pattern and mitonuclear co-introgression test

A genome-wide scan using 50 kb non-overlapping windows for four detected hybridization events revealed distinct introgression landscapes; however, all were characterized by introgression islands, primarily concentrated at the distal ends of smaller chromosomes beyond chromosome 11. The *S. maurus*–*S. stejnegeri* pair exhibited the most introgression islands, followed by *S. dacotiae*–*S. torquatus* and *S. maurus*–*S. rubicola*, while *S. maurus*–*S. dacotiae* showed few islands. Pairwise correlation analyses revealed positive correlations (*P* < 0.01) for all pairwise comparisons, except between *S. maurus*–*S. dacotiae* and *S. dacotiae*–*S. torquatus*, which was not statistically significant (*P* = 0.35). Notably, the highest correlation coefficient (ρ = 0.44) was found between *S. maurus*–*S. dacotiae* and *S. maurus*–*S. rubicola*, further suggesting that introgression between *S. maurus* and *S. rubicola*/*S. dacotiae* may have occurred in the MRCA of the latter two species.

Using a threshold of *f*_d_ = 0.2, 894–2,275 introgressed genes were identified across four species pairs. GO enrichment analysis revealed significant functional enrichments in three species pairs (*P*_adjust_<0.05): 1,323 genes between *S. maurus* and *S. rubicola* were enriched in phosphatidylethanolamine metabolic process (GO:0046337); 2,275 genes between S*. maurus* and *S. stejnegeri* in intermediate filament organization (GO:0045109); and 2,093 genes between *S. dacotiae* and *S. torquatus* in ADP-ribosyltransferase (GO:0106274, GO:1990404, GO:0003953) and pentosyltransferase activity (GO:0016763) (Figure S6). KEGG analysis showed significant enrichment only for *S. maurus*–*S. stejnegeri* genes in metabolism of xenobiotics by cytochrome P450 (KO00980), while other pairs exhibited no significant enrichment (*P*_adjust_>0.05) (Figure S7). Notably, in the GO enrichment results for *S. maurus*–*S. rubicola* introgressed genes, there is a term mitochondrial translation (GO:0032543) ranked third by adjusted *P* value, though statistically non-significant (*P*_adjus_=0.26). This term encompasses mitochondrial ribosomal (e.g., *MRPS16*, *MRPL23*), tRNA-related (e.g., *SARS2*, *TRMT10C*), and translational regulator (e.g., *MTIF3*, *TSFM*) genes, which possibly indicate mitonuclear co-introgression between *S. maurus* and *S. rubicola* or between *S. maurus* and MRCA of *S. rubicola* and *S. dacotiae* (or its sister species).

Using various methods for introgression gene detection, along with different introgression indicator, thresholds and species combinations, nearly half of the mitonuclear co-introgression tests yielded statistically significant results (*P*<0.05; Tables S3, S4). Considering that *S. rubicola* and *S. dacotiae* may not fully represent their ancestors or its sister lineage, these results may support our hypothesis that the cytonuclear conflict of *S. maurus* originates from mitogenome capture from the MRCA of *S. rubicola* and *S. dacotiae* (or its sister species), accompanied by mitonuclear co-introgression. Notably, majority of significant results were observed when *O. deserti* was used as the outgroup, whereas tests using *S. stejnegeri* or *S. rubetra* as the outgroup were largely non-significant. This may be because these two species are too closely related to the target species (or have hybridized with it) and thus cannot reliably represent ancestral alleles.

### Demographic history over the past million years

Over the past million years, four *Saxicola* species have undergone significant population fluctuations in their demographic histories. The range *Ne* fluctuations vary greatly among the different species. The maximum *Ne* of *S. rubetra* reached 8-9 million individuals, while that of *S. rubicola* was less than 90,000 individuals, differing by two orders of magnitude. The maximum *Ne* of *S. maurus* and *S. stejnegeri* are generally similar, around 1-2 million individuals.

The *S. rubetra* population began to grow earliest, starting from ∼1 Mya, with a plateau period from ∼0.4 Mya to ∼0.2 Mya, followed by further growth starting around 0.2 Mya. The most significant growth occurred around 0.1 Mya (just before or at the onset of the Last Glacial Period), followed by a rapid collapse around 0.07 Mya. *S. stejnegeri* and *S. maurus* showed comparable growth starting around 0.4–0.3 Mya, but S*. maurus* expanded more rapidly, exhibiting a sharp increase around 0.05–0.04 Mya followed by a dramatic contraction around 0.03–0.02 Mya, coinciding with the Last Glacial Maximum. *S. rubicola* exhibited a particularly distinctive demographic trend: following a slight increase around 1 Mya, it underwent a sharp decline, then rebounded around 0.2 Mya, coinciding with the onset of the Last Glacial Period, and returned to its original level by ∼0.04 Mya before declining again—forming a wave-like pattern of population fluctuation.

## Discussion

### Maternal co-inheritance and discordance with species tree

Previous phylogenies based on single mitochondrial genes suggested that *S. maurus* is sister to *S. dacotiae* and *S. rubicola*, relative to *S. torquatus* (Zink et al. 2009). Our mitogenome-based phylogeny further supports this relationship. We also found that the NRW tree aligns with the mitogenome tree but diverges from the species tree, indicating maternal coinheritance. Similar cases have been reported in other avian lineages, such as *Ficedula* (Smeds et al. 2015), *Anas* (Gu et al. 2023), and *Emberiza* (Zhang et al. 2024). Though paternal leakage of mtDNA or recombination in either molecule could potentially weaken the evolutionary association between the NRW and the mitogenome (White et al. 2008). Nevertheless, our findings stress that the NRW and mitogenome exhibit congruent lineage characteristics, and their evolutionary stability in stonechat aligns with mainstream opinions (Ellegren 2007).

One possible perspective is that the mitogenome and NRW represent the true species tree, as these two molecular markers do not recombine and can therefore reflect their true evolutionary history (Moore 1995, Mackiewicz et al. 2019). In contrast, nuclear markers may have undergone genome homogenization due to introgression between *S. torquatus* and *S. rubicola*/*S. dacotiae*, or between *S. maurus* and *S. stejnegeri*, leading to shared genetic material between corresponding species pairs. This would bring the species pairs closer together on phylogenetic trees based on nuclear SNVs. However, this scenario should result in substantial topological conflicts at corresponding nodes (Steenwyk et al. 2023). Our result rejects the mitogenome and NRW tree as the species tree (Figure 2 a). Similarly, the very low proportion (∼1.7%) of the topology consistent with the mitogenome/NRW tree in the GF trees further rejects this possibility as the species tree (Figure 1 c).

We found eight hypotheses that could explain the incongruence between mtDNA/NRW tree and species tree (Figure 5 a-h). Considering the introgression and mitonuclear co-introgression signals detected between *S. maurus* and MRCA of *S. rubicola* and *S. dacotiae*, as well as their proximity in the breeding range, the incongruence may result from the mitogenome/W chromosome of *S. mauru*s being captured from the MRCA of *S. dacotiae* and *S. rubicola* (Figure 5 a). However, due to incomplete sampling within the breeding ranges of *S. dacotiae* and *S. rubicola*, several alternative capture scenarios cannot be mutually exclusive (Figure 5 b-e). Except for ILS-generated lineages (Figure 5 e), other scenarios (Figure 5 b-d) could yield observed introgression and mitonuclear co-introgression signals between *S. maurus* and MRCA of *S. rubicola* and *S. dacotiae*, as the other mitogenome donor lineages share genetic similarity with this MRCA. However, the fourth hypothesis (Figure 5 d) is less probable: female-specific selection strong enough to create additional maternal lineages without speciation (e.g., nest parasitism) (Fossøy et al. 2016) has not been observed in stonechat, and high male-biased dispersal leading to additional lineages is unlikely in stonechat, especially in songbirds, which typically exhibit female-biased dispersal (Clarke et al. 1997). The fifth and sixth hypothesis (Figure 5 e, f) is also improbable given low detected ILS signals (Fig. 2 b) and long, well-supported intermediate branches in mitogenome ML trees (Figure 1 a), which are inconsistent with ILS expectations (Avise and Robinson 2008, Rivas-González et al. 2023). The two latest hypotheses (Figure 5 g, h) are also less probable, as no introgression signals exist between *S. torquatus* and lineages genetically similar to this MRCA (e.g., *S. stejnegeri*/*S. maurus*). Therefore, overall, compared to the other hypotheses, the first three hypotheses (Figure 5 a, b, c) are the most likely, i.e., the mitogenome/W chromosome of *S. maurus* may have been captured from lineage related to the MRCA of *S. dacotiae* and *S. rubicola*. Future population genomic analysis with extensive sampling in breeding areas will further validate this and may distinguish the second hypothesis from the other two.

**Figure 5.**
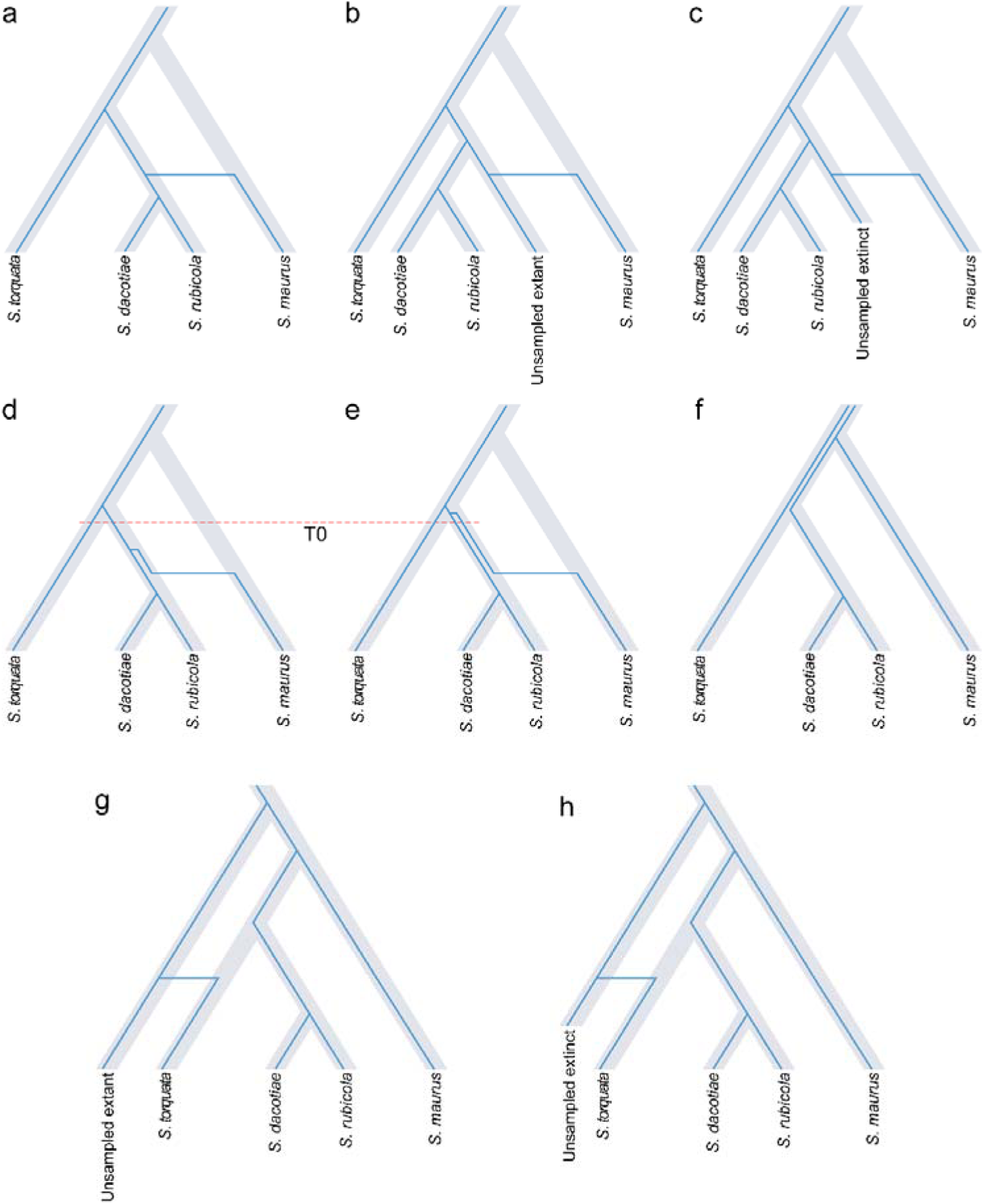
Eight hypotheses potentially explaining the observed incongruence between the species tree and mitogenome/NRW trees. The incongruence may result from the mitogenome/W chromosome of *S. maurus* being captured from: (a) the MRCA of *S. dacotiae* and *S. rubicola*; (b) an unsampled extant lineage sister to the MRCA of *S. dacotiae* and *S. rubicola*; (c) an extinct lineage sister to the MRCA of *S. dacotiae* and *S. rubicola*; (d) an unsampled lineage within the MRCA of *S. dacotiae* and *S. rubicola* generated by female-specific selection or male-biased dispersal; (e) an unsampled lineage within the MRCA of *S. dacotiae* and *S. rubicola* generated by ILS from ancestral polymorphisms in the MRCA of *S. torquatus*, *S. dacotiae*, and *S. rubicola*. The incongruence may have arisen directly from ILS resulting from ancestral polymorphisms in the MRCA of *S. torquatus*, *S. dacotiae*, *S. rubicola*, and *S. maurus* (f). The mitogenome/W chromosome of *S. torquatus* may have been captured from an unsampled extant (g) or extinct (h) lineage that is sister to the MRCA of *S. dacotiae*, *S. rubicola*, *S. maurus* and *S. torquatus*. The species tree is shown in grey with the blue lines representing mitogenome/W chromosome lineages. The divergence time (T0) between different lineages of mitogenome/W chromosome within the MRCA of *S. dacotiae* and *S. rubicola* would be shallower than the root (d) or around the root divergence time (e).

### Plausibility of introgression within the *S. torquatus* complex

Species undergoing rapid radiation are prone to hybridization due to frequent geographic overlap with closely related taxa (Edelman et al. 2019). We reveal extensive hybridization among several species in the *S. torquatus* complex, which is plausible when considering the distribution of their breeding ranges. For example, *S. maurus* and *S. stejnegeri* exhibit parapatric distribution along the Yenisei/Angara River, where hybrids between them have been reported (Hellström and Norevik 2014, Hellström and Waern 2023). Similarly, *S. dacotiae* and *S. torquatus* may come into contact in northwestern Africa, as *S. torquatus* has scattered populations in the northern part of the continent. Additionally, *S. maurus* could have hybridized with the MRCA of *S. dacotiae* and *S. rubicola* when considering that *S. dacotiae* and *S. rubicola* may gradually colonize Europe from West Asia (Illera et al. 2008).

Moreover, we identified hybridization between *S. maurus* and the MRCA of *S. rubicola* and *S. dacotiae* via *f_b_*, which was further supported by the high correlation of introgressed regions between *S. maurus*–*S. rubicola* and *S. maurus*–*S. dacotiae*. However, there was a notable discrepancy: a strong introgression signal between *S. maurus* and *S. rubicola*, but a weak signal between *S. maurus* and *S. dacotiae*. This discrepancy may be due to additional introgression between *S. maurus* and *S. rubicola*. Another possibility is a limitation of the ABBA-BABA test (The Heliconius Genome Consortium 2012): since introgression between *S. torquatus* and *S. dacotiae* was detected, using the (((*S. torquatus*, *S. dacotiae*), *S. maurus*), Outgroup) combination to test for introgression between *S. dacotiae* and *S. maurus* might have obscured the signal due to introgression between *S. torquatus* and *S. dacotiae*. The third potential reason is that the island species *S. dacotiae*, with a small population size, has accumulated more derived mutations due to larger genetic drift, which may lead to the loss of ancestral introgression signals. This is supported by the relatively long terminal branch lengths in genome-wide SNVs and mitogenome ML trees (Figure 1 a), and previous studies suggest that differentiation of *S. dacotiae* from other *Saxicola* species has largely been driven by genetic drift (Van Doren et al. 2017). Similarly, in the Phylonet species network, one of the hybridization events, where introgression between *S. maurus* and *S. rubicola*, rather than between *S. maurus* and the MRCA of *S. rubicola* and *S. dacotiae*, could also be influenced by these factors. Furthermore, Treemix identified hybridization events involving *S. rubetra* with *S. torquatus* and *S. rubicola*. As these events were not detected by other methods, they may represent spurious introgression signals due to sampling gaps of intermediate evolutionary lineages or larger fluctuations in allele frequencies due to small sample sizes per species (Pickrell and Pritchard 2012), despite past reports of hybridization between *S. rubetra* and *S. torquatus* complex species (McCarthy 2006).

### Drivers of genomic introgression landscapes

The genomic landscape of introgression is typically influenced by the interplay between recombination rates and selection against genomic incompatibilities (Martin et al. 2019, Duranton and Pool 2022, Veller et al. 2023). When selection against genomic incompatibilities is the dominant force, introgression is expected to exhibit a positive correlation with recombination rate (Martin et al. 2019, Edelman et al. 2019); conversely, if foreign ancestry is favored, a negative correlation is anticipated (Duranton and Pool 2022).

We found that strong introgression regions tend to be concentrated in island-like forms at the ends of small chromosomes, likely due to the higher recombination rates in these regions (Ellegren 2013, Zhang et al. 2014, Guerrero-Cózar et al. 2021, Leitwein et al. 2024), although the lack of high-resolution recombination maps limits further validation. A similar pattern has been observed in other groups. For example, in Princess cichlids, introgressed segments are significantly enriched at the chromosome periphery, consistent with their higher recombination rates compared to the central regions (Gante et al. 2016, Leitwein et al. 2024). Similarly, in Amazonian antbird (*Thamnophilus aethiops*) (Musher et al. 2024) and Swainson’s thrushes (*Catharus ustulatus*) (Blain et al. 2025), introgression signals are mainly concentrated on small chromosomes, which aligns with their higher recombination rates relative to large chromosomes (Ellegren 2013, Zhang et al. 2014).

This enrichment pattern of introgression in regions with potentially higher recombination rates also suggests that in the *Saxicola* clade, foreign ancestry may have been subject to strong purifying selection due to genetic incompatibilities, consistent with general observations (Presgraves 2010, Moran et al. 2021). However, functional enrichments of introgressed genes were also detected in multiple species pairs, including pathways like phospholipid metabolism, cytoskeleton organization, and detoxification metabolism, indicating the presence of adaptive introgression. Therefore, while not excluding other factors, the introgression landscape in several *Saxicola* species is likely shaped by the interplay between strong selection against genomic incompatibilities and weak adaptive introgression.

Moreover, correlation of introgression between different species is always positive, even when ancestral introgression is not considered. Given that introgressed genes are not functionally enriched in similar pathways across species pairs, this correlation may mainly stem from intrinsic properties of specific genomic regions (e.g., shared high-recombination areas) rather than parallel selection acting on particular functional candidate genes.

## Conclusion

Our genome-wide analyses encompassed all species-level lineages within the *S. torquatus* complex, except *S. tectes*. We recovered robust phylogeny and revealed widespread phylogenetic discordance across genomic regions, primarily driven by introgression rather than ILS, showing reticulate evolutionary history. Cytonuclear discordance likely resulted from introgression and reflects maternal co-inheritance of mitogenome and the W chromosome. The Z chromosome exhibited greater phylogenetic stability, being less affected by ILS and introgression, showing its robustness in species tree reconstruction (also see in Zhang et al. 2024, 2025b, Stiller et al. 2024). Introgressed regions were concentrated at the distal ends of smaller chromosomes and correlated across species pairs, likely attributed to shared recombination hotspots in these regions. Some implications can be drawn from these results. First, our phylogenomic analysis positions *S. stejnegeri* as the most basal lineage within the *S. torquatus* complex, whereas *S. maurus* is sister to all other members. Together with acoustic evidence (Opaev et al. 2018), this study supports recognizing *S. stejnegeri* as a distinct species. Additionally, the new phylogenetic relationship, where *S. stejnegeri* and *S. maurus* are sequentially nested at the root, further reinforces the Asian origin of this complex (Figure 6) (Illera et al. 2008, Jiang et al. 2025). Although the precise ancestral ranges of *S. stejnegeri* and *S. maurus* in Siberia remain uncertain, European and African populations likely originated from later colonization by *S. maurus* ancestors. Furthermore, genomic evidence of introgression between the ancestral European lineage (MRCA of *S. rubicola* and *S. dacotiae*) and *S. maurus* favors an Asian (rather than African) origin for European population, followed by Canary Islands colonization. African populations may have been colonized either through Europe or via a separate dispersal directly from Asia, followed by colonization to Réunion Island (Illera et al. 2008, Jiang et al. 2025). Future genome-wide analyses with comprehensive sampling across *Saxicola* will provide deeper insights into their colonization history.

**Figure 6.**
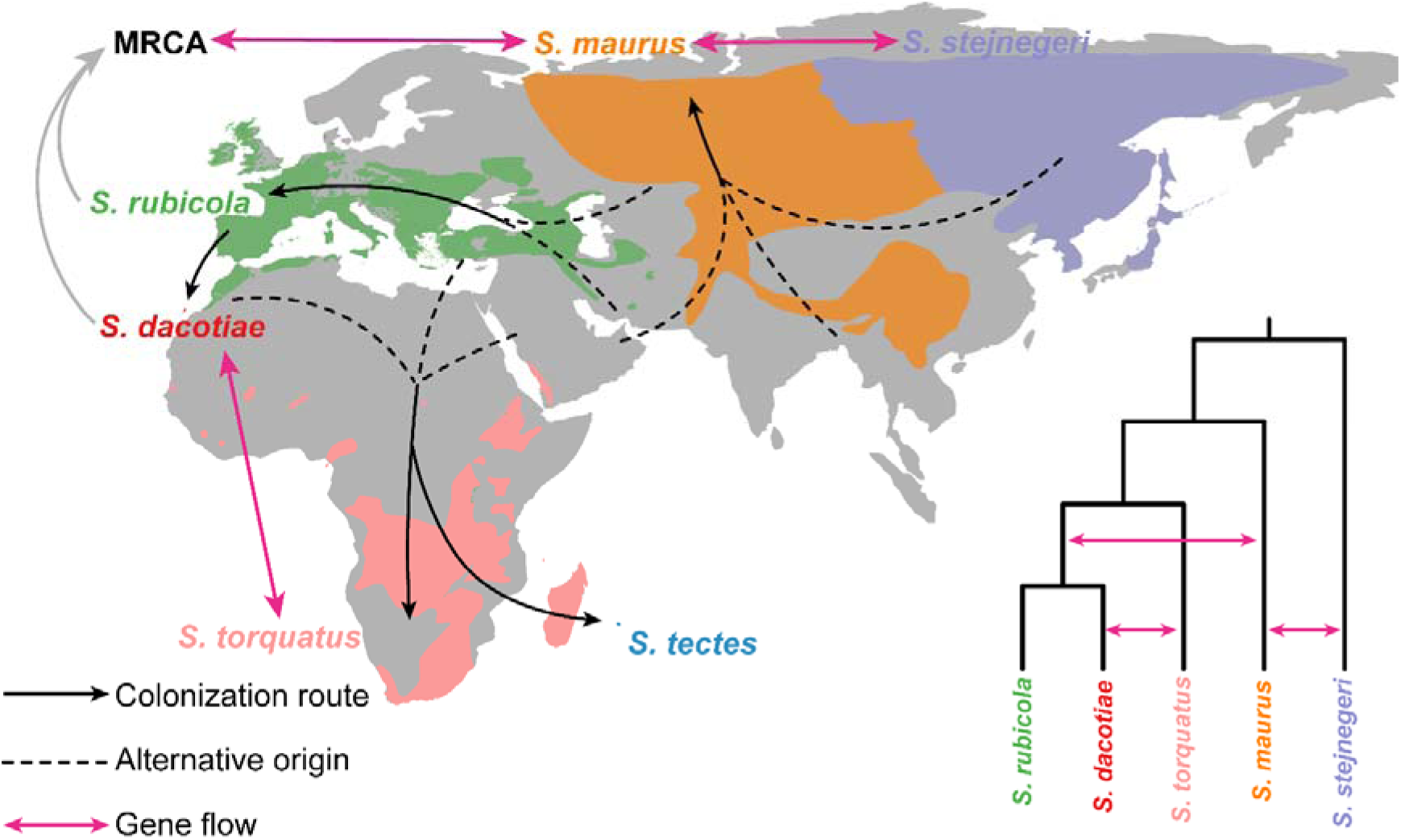
Schematic diagram of colonization and hybridization in five *S. torquatus* complex species. Colored regions denote the breeding areas of each species, based on data from Birds of the World (Billerman et al. 2022). The breeding ranges of *S. maurus* and *S. stejnegeri* are separated by the Yenisey/Angara rivers. Black arrows indicate colonization routes, with dashed lines at the starting points representing alternative origins. A gray arrow points from *S. rubicola* and *S. dacotiae* to their MRCA. Red bidirectional arrows on both the map and the phylogenetic tree denote gene flow.

## Author contributions

C.J. and B.L. designed the research and collected the samples. C.J. performed the experiments and data analysis. W. Z. and Z. J. contributed to data analysis. C. J., Y. L., and W. Z. wrote the manuscript. N.M. and B.L. revised the manuscript and improved the language. All authors have read and agreed to the final version of the manuscript.

## Acknowledgments

We are grateful to Wei Zhang, Yanchun Xu, and Suying Bai for their guidance and logistical support during sample collection.

