## Supplementary figures and images for "Genome-wide SNVs revealed reticulate evolutionary history of an Eurasia-African songbird radiation"

### Fig S1.jpg

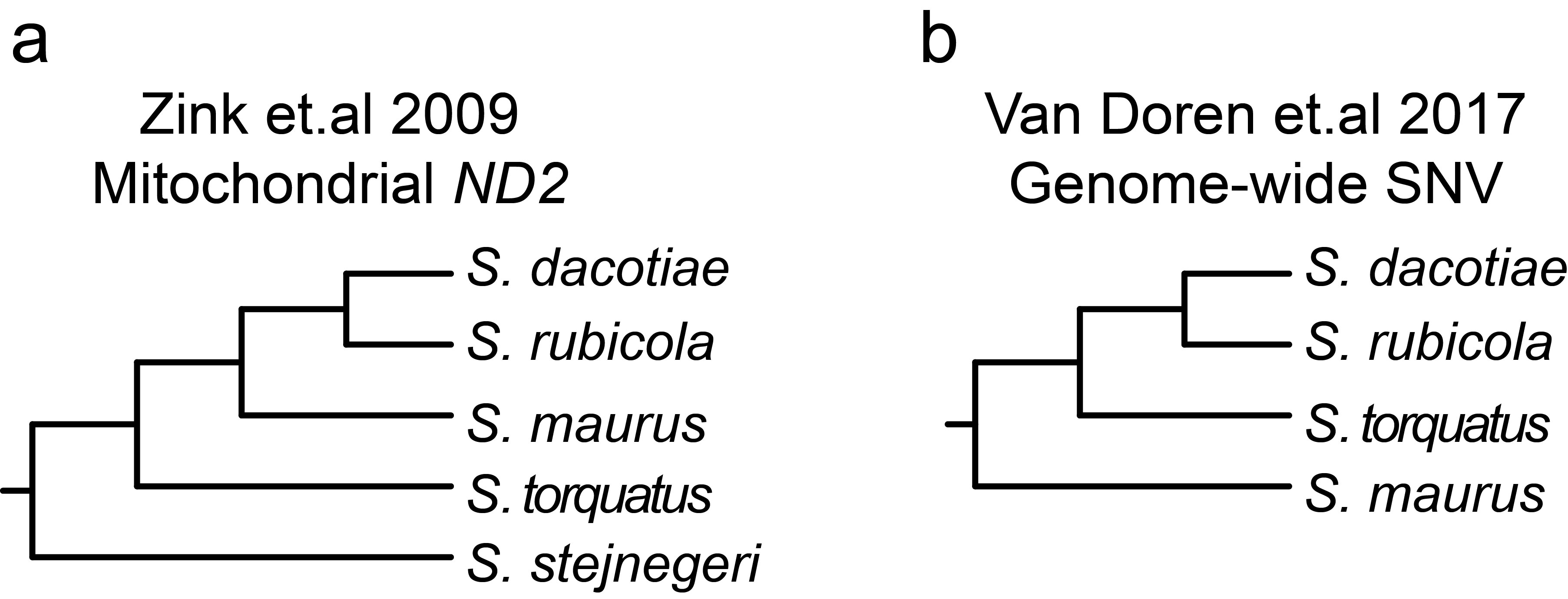

### Fig S2.jpg

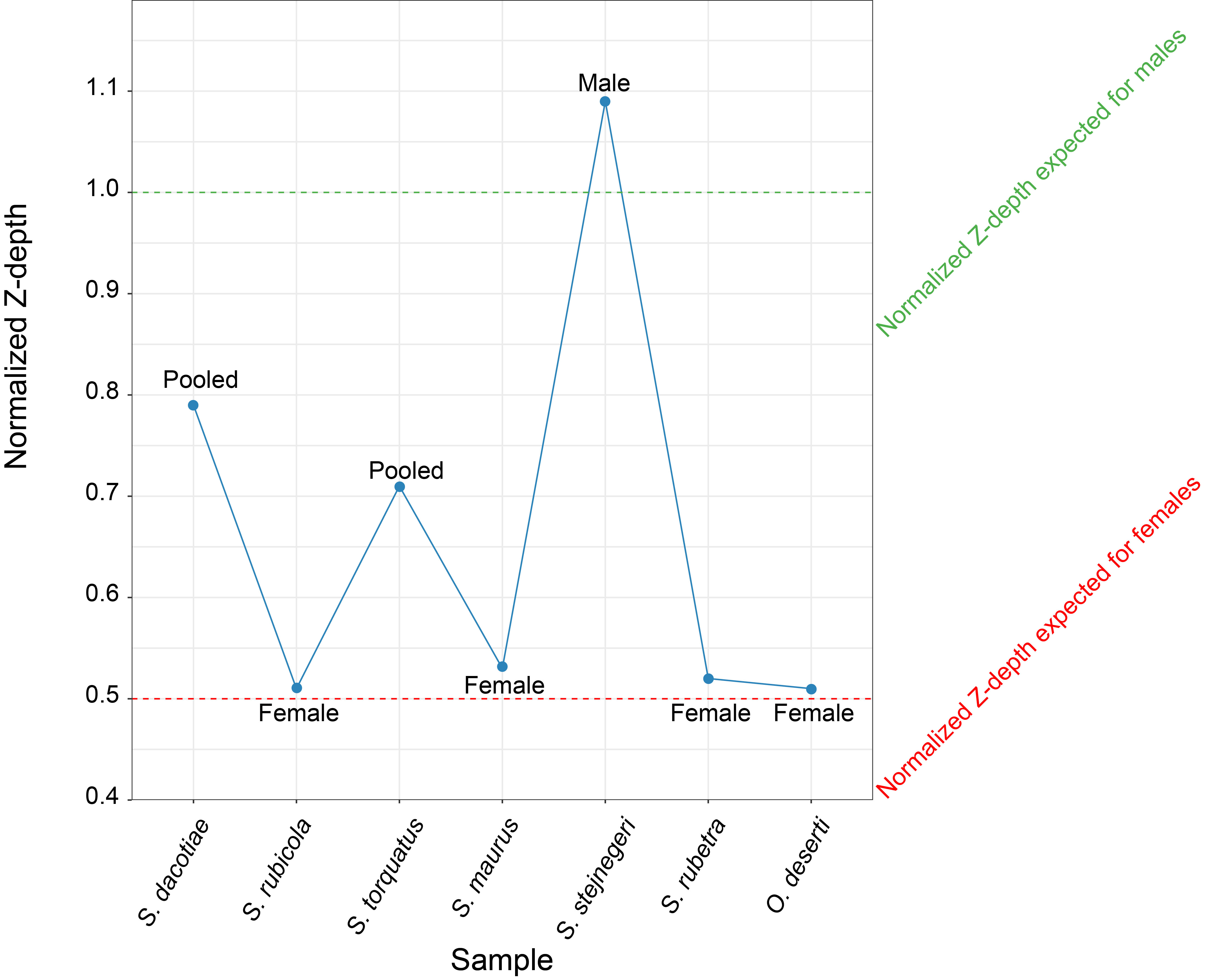

### Fig S3.jpg

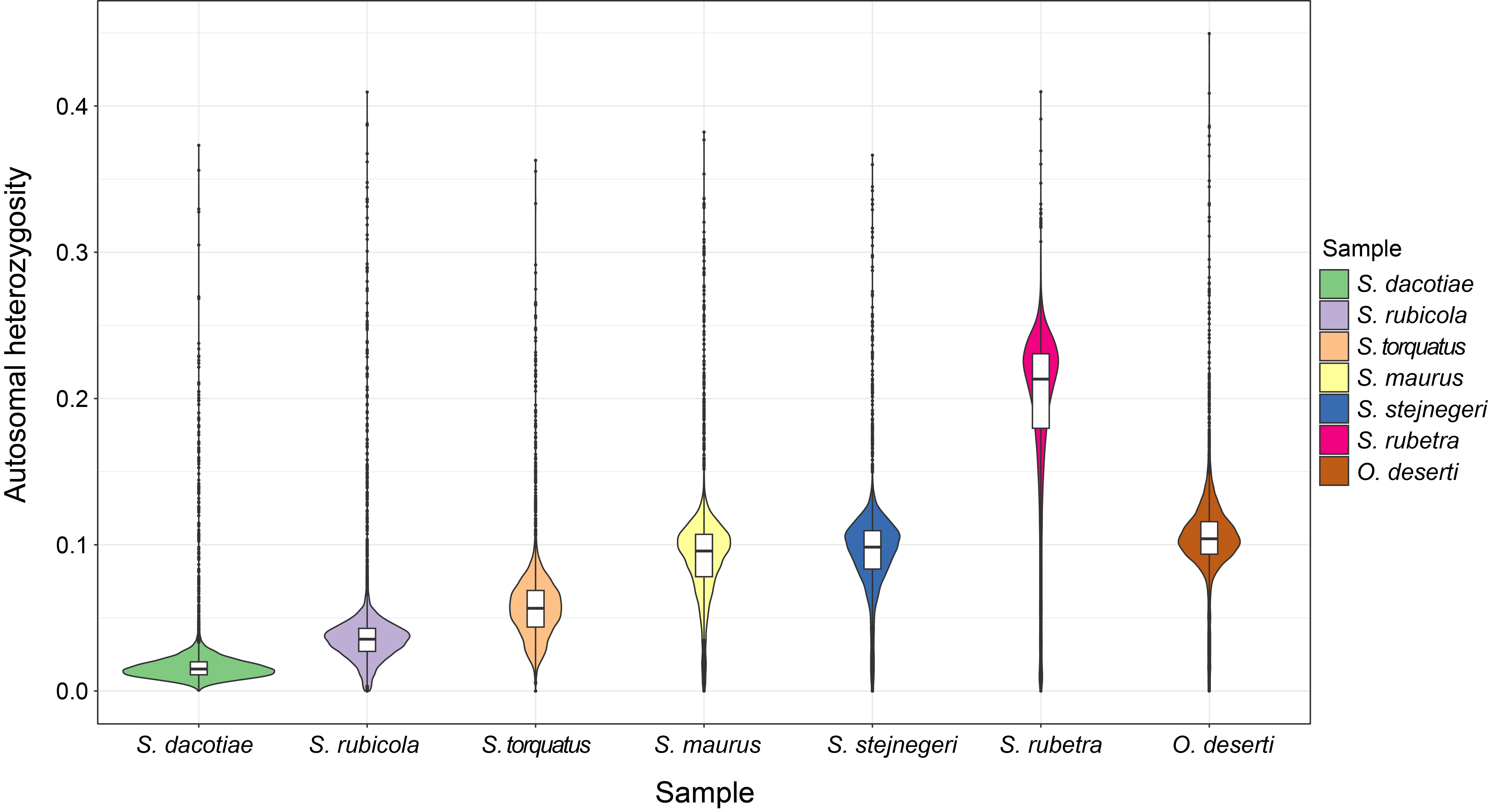

### Fig S4.jpg

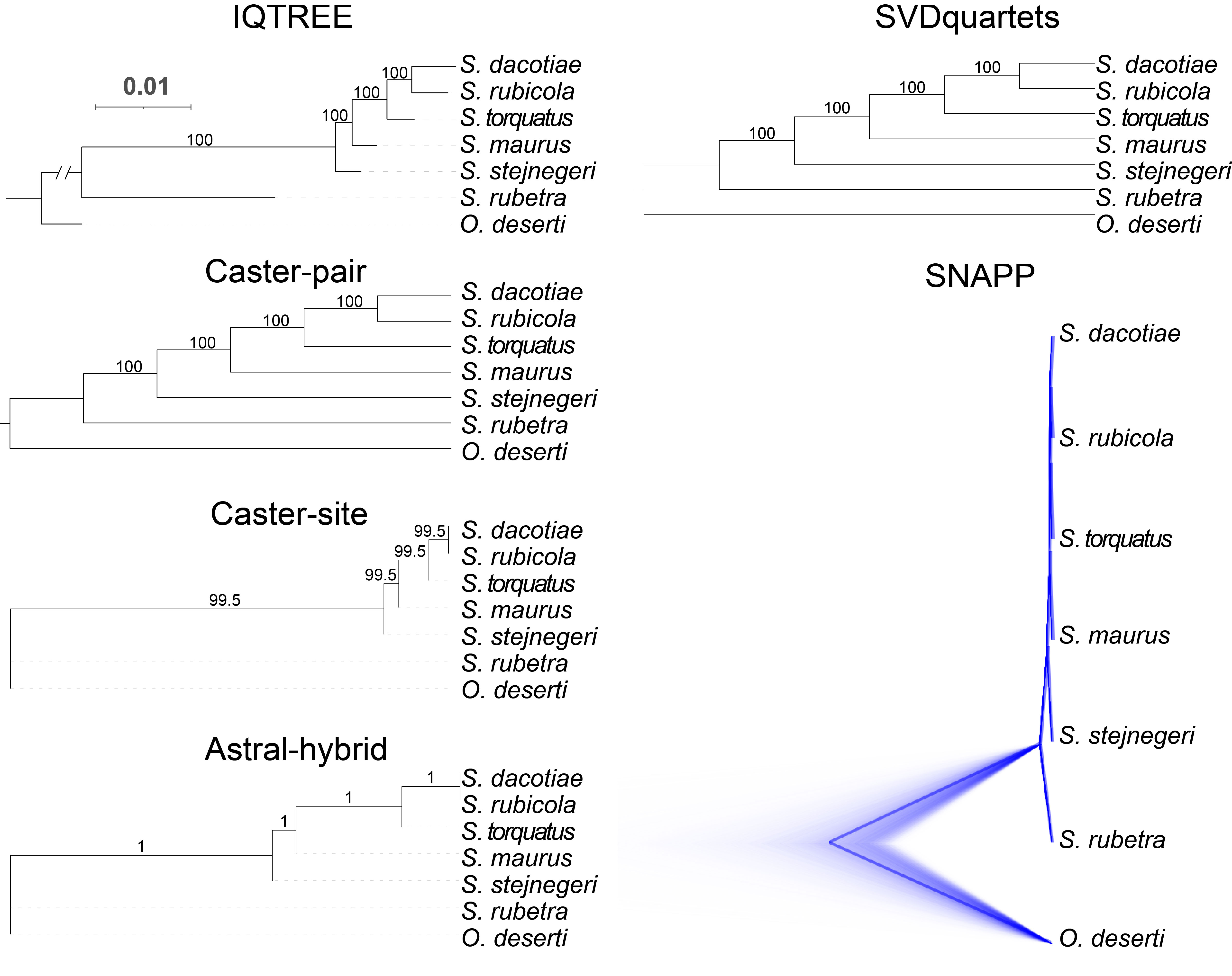

### Fig S5.jpg

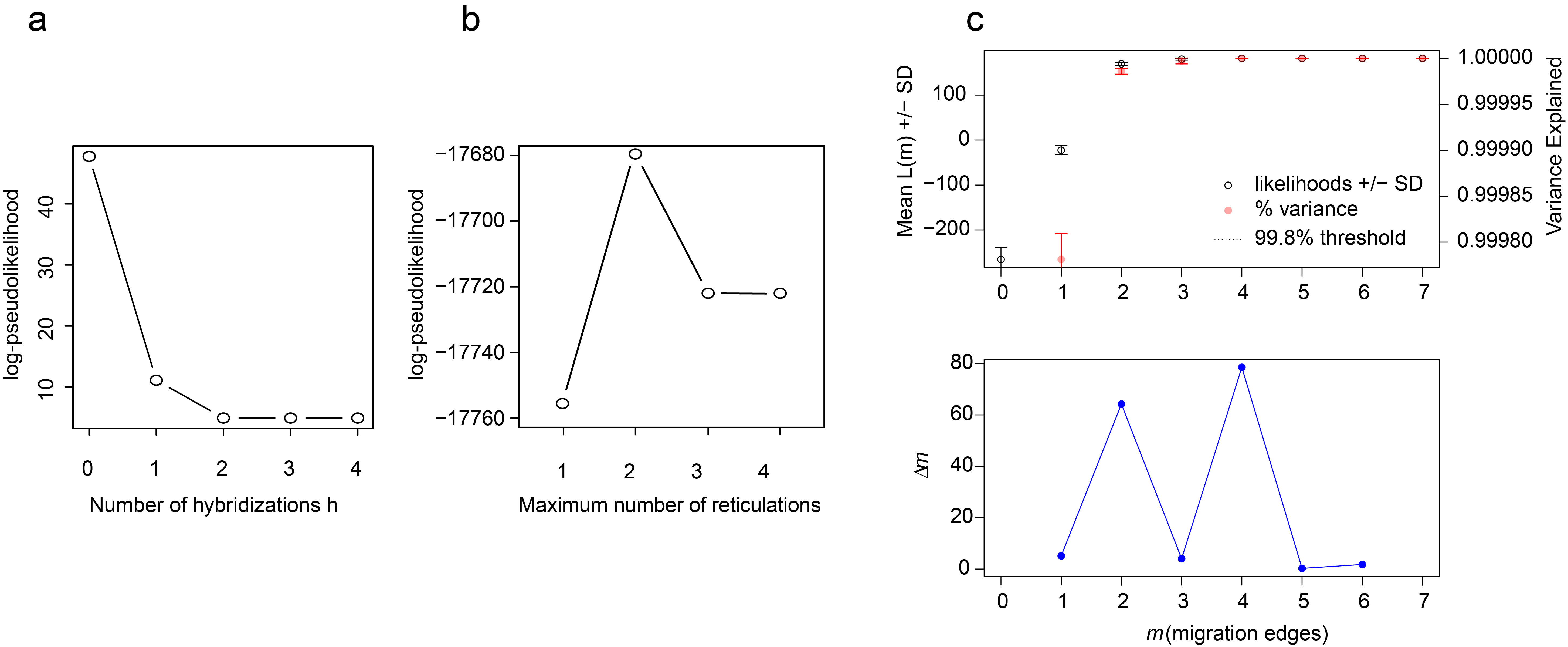

### Fig S6.jpg

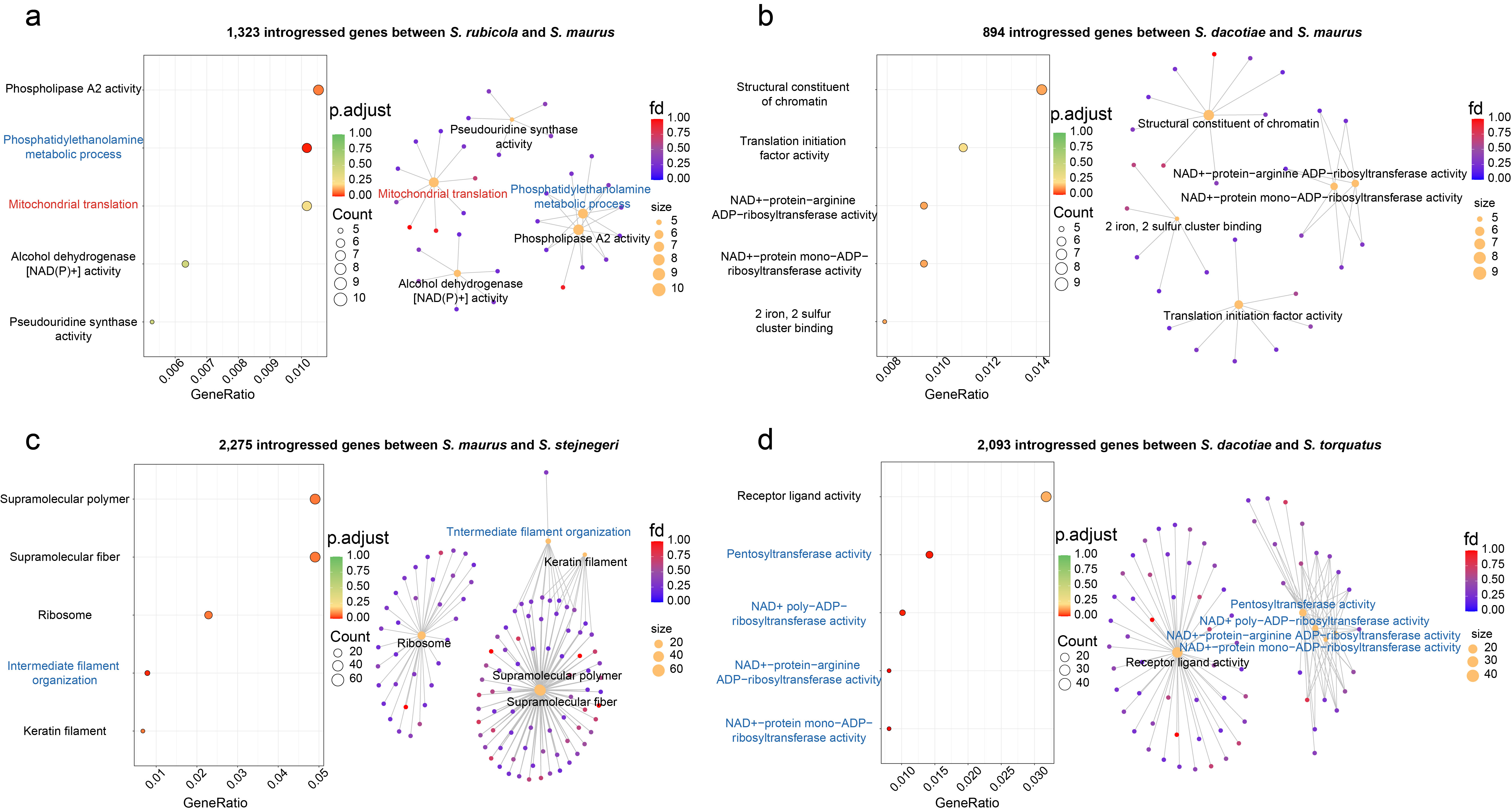

### Fig S7.jpg

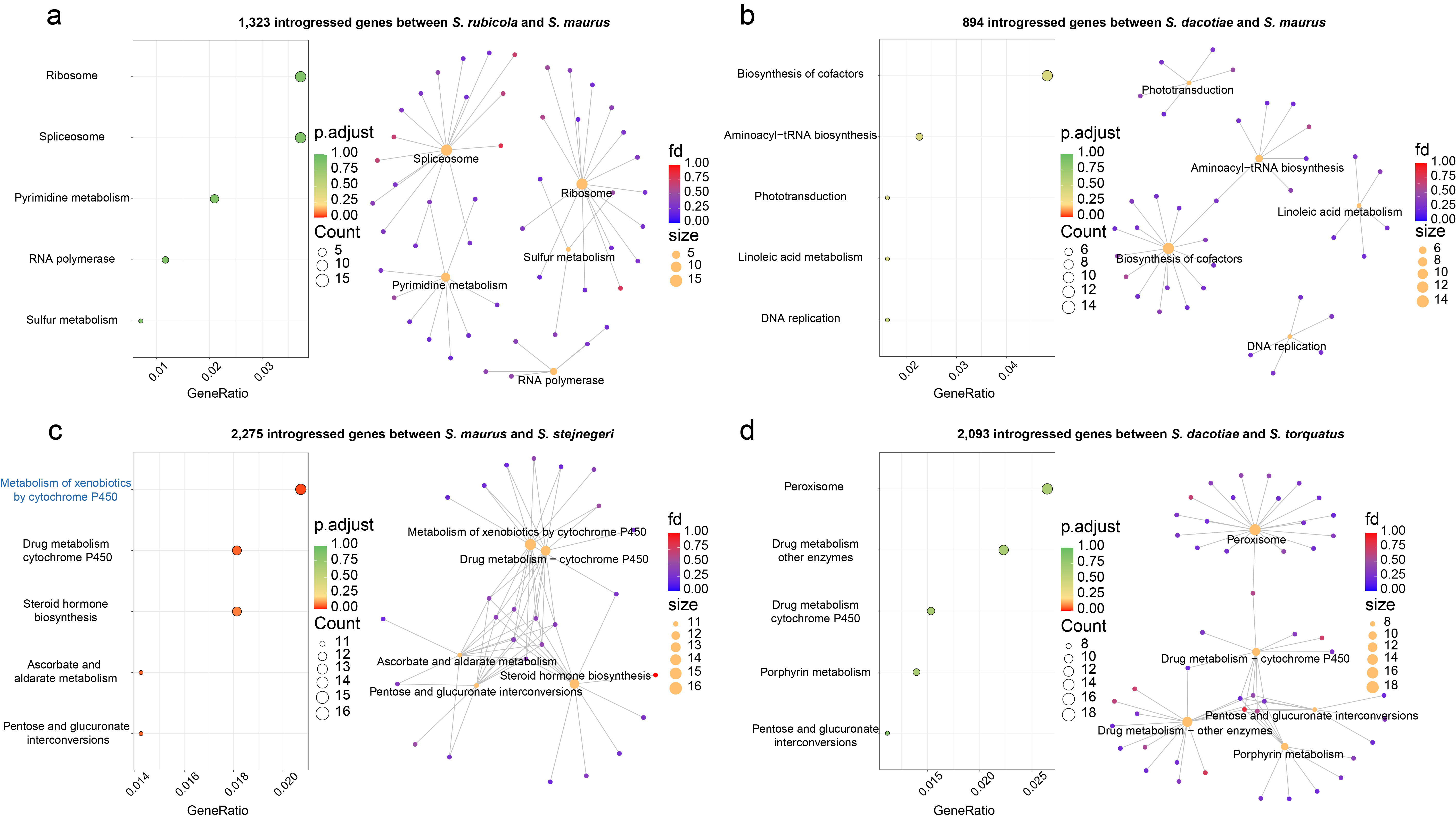
